# Diatom Endosymbionts have Shrinking but Stable Genomes Despite Low Coding Density

**DOI:** 10.64898/2026.03.19.712447

**Authors:** Heidi Abresch, Sophia Miller, Forest Cruse, Jingchun Li, Sarah Hamsher, J. Patrick Kociolek, Scott R. Miller

## Abstract

Successful establishment of long-term, obligate endosymbiotic relationships requires integration of hosts and endosymbionts across multiple levels. For example, highly integrated, host-beneficial endosymbionts typically have extremely reduced genomes and metabolisms. However, we do not yet fully understand the specific mechanisms that drive this integration or if there is a specific order in which these changes must occur. To investigate the early stages of endosymbiont genome reduction, we greatly expanded available whole genome data for the nitrogen-fixing endosymbionts (spheroid bodies, SBs) of diatoms in the family Rhopalodiaceae. We used these data to reconstruct SB evolutionary history and to characterize SB core metabolic capacity. We found two key genes missing from all SB genomes, *mltA* and *dnaA*, which could provide points of host control over SB cell division. Although most of the SB core genome is experiencing moderately strong purifying selection, we identified 54 genes under positive selection. Eighteen of these are peripheral proteins or involved in cell wall and cell membrane metabolism and could be involved in direct interactions with the host. Unexpectedly, we also found three *nif* genes under positive selection that are core to the central nitrogen-fixing enzyme. Overall, our results provide early insights into how SBs and their hosts interact, showing that SBs are still in the early stages of endosymbiont genome reduction, but they differ in key ways from current models, including the early loss of all mobile elements.

## Introduction

Host-microbe endosymbioses are ubiquitous in nature and have massive impacts on the ecology and evolution of both partners (Bennett and Moran, 2015; Husnik et al., 2021). While many of these relationships remain facultative, some become obligate, where the two partners can no longer survive independently. Successful establishment of these long-term, obligate endosymbiotic relationships requires integration of hosts and endosymbionts on genetic, metabolic, and cellular levels. For example, highly derived endosymbionts typically have extraordinarily reduced genomes with specialized metabolic capacity, rely on imported host-derived proteins, and divide synchronously with their hosts (McCutcheon and Moran, 2012).

Although there are many examples of how tight-knit endosymbiotic relationships look after hundreds of millions or even greater than a billion years of evolution, we do not fully understand the specific mechanisms that drive this integration, or if there is a particular order in which these changes need to occur. Addressing the mechanisms of early integration between partners is crucial to explain how long-lasting endosymbiotic relationships successfully establish. Furthermore, most models of endosymbiont evolution are systems with invertebrate hosts (McCutcheon and Moran, 2012; McCutcheon, 2021), possibly obscuring which features of endosymbiont evolution are universal and which features are a result of different host contexts (Husnik and Keeling, 2019; Husnik et al., 2021). To address these outstanding questions, we need more diverse sampling of more recent endosymbiotic associations, particularly from single-celled hosts.

One emerging system for studying endosymbiont evolution is between nitrogen (N)-fixing endosymbionts called spheroid bodies (SBs), which are derived from a unicellular bacterium, and their diatom hosts (a clade of unicellular photosynthetic stramenopiles) in the family Rhopalodiaceae (Nakayama et al., 2014). This system is one of the most geographically widespread eukaryote N-fixing associations (Schvarcz et al., 2022), contributing significantly to the global N cycle.

There are two key reasons SBs are a promising model system to study early endosymbiont evolution. First, SBs are relatively young endosymbionts (at least 34 Mya; (Nakayama et al., 2014)) compared with well-studied insect-endosymbiont systems (typically *>*100 to over 270 Mya; (McCutcheon, 2021)), yet SBs exhibit important signs of host-integration, including highly altered metabolisms. For example, they rely on their hosts for carbon, as they have lost most genes necessary for chlorophyll synthesis and carbon fixation (Nakayama and Inagaki, 2017; Abresch et al., 2024). There is also evidence for cellular integration, as SBs are passed down uniparentally during sexual reproduction in *Rhopalodia gibba* (Kamakura et al., 2021). Second, SBs are closely related to the recently described N-fixing endosymbiotic organelle, the nitroplast (previously identified as UCYN-A2; (Abresch et al., 2024; Coale et al., 2024)). SBs and nitroplasts share intriguing similarities: 1) they both fix N for their hosts in exchange for fixed carbon (Coale et al., 2024; Marks et al., 2025); 2) they live in physiologically similar host environments (single-celled, photosynthetic hosts); 3) their hosts—at least ancestrally—occupied similar niches (oligotrophic ocean). However, the nitroplast is estimated to be an older endosymbiotic relationship, the host provides targeted protein import, and cell division is tightly coordinated with the host (Coale et al., 2024). The close evolutionary relationship and functional similarities between SBs and UCYN-A means SBs could provide a possible proxy for the earlier stages of nitroplast evolution before becoming fully integrated with their hosts.

However, critical questions about SBs are unresolved, including their position along the trajectory of endosymbiont evolution and to what extent they are integrated with their hosts. Although SB genomes are about 60% smaller than free-living relatives (Abresch et al., 2024), they are not yet as highly reduced as those from the best studied endosymbiont systems. This allows for a deeper examination of how genes are lost over time during endosymbiotic evolution. Previous work also suggests that SBs may not fully align with the current model of endosymbiont genome evolution, which proposes that transposases drive early genome reduction (McCutcheon, 2021); however the first four analyzed SB genomes completely lack selfish elements (Abresch et al., 2024). There are also conflicting results regarding the identity of the closest free-living relative of SBs, which muddies our understanding of SB evolutionary origins (Schvarcz et al., 2022; Abresch et al., 2024). Finally, we don’t know how SBs and hosts directly interact or exchange nutrients. A larger dataset of SB genomes from diverse hosts would allow us to explore these critical aspects of the SB system and address broader evolutionary questions about endosymbiont evolution. By identifying the SB core genome and metabolic capacity, we can identify the absence of specific “essential” genes from SB genomes and highlight possible points of host control over SBs. Furthermore, points of direct interaction between host and SBs may be revealed by signatures of positive selection (Bennett and Moran, 2015).

To address these outstanding questions, we cultured diverse SB-hosting diatom taxa from a variety of locations and assembed 22 new SB genomes—a three-fold increase in the number of available SB genomes. First, we compared general patterns of SB genome characteristics to current models of endosymbiont genome reduction. Second, we determined the closest free-living relatives of SBs and relationships among SBs from different host taxa by constructing a genome-wide phylogeny. Third, we characterize SB core metabolic capacity and possible mechanisms of host control over SB growth and division by estimating the SB core genome. Finally, we identified SB genes that putatively interact with the host directly by analyzing the strength and direction of selection on core genes. Overall, our results show that SBs are still in the early stages of endosymbiont evolution, but they differ in key ways from current models of endosymbiont genome reduction. Specifically, these results can guide further investigations of how the diatom hosts and SBs are genetically and metabolically integrated. Broadly, this study provides fresh insights into the general mechanisms of endosymbiont genome reduction and helps identify key processes in the early stages of endosymbiont evolution in unicellular hosts.

## Results and Discussion

### Spheroid body genomes have low coding density but high synteny

Laboratory strains of multiple Rhopalodiaceae species were cultured from samples collected between August 2020 and May 2023 from 15 locations across 6 US states (Figure 1, Supplemental Table 1). From these, we assembled 22 new SB genomes from short-read sequence data. SB assemblies ranged from closed chromosomes to 477 contigs; the draft assemblies were essentially complete in terms of gene content based on comparable CheckM scores with the closed SB genomes (Supplemental Table 1).

**Fig. 1:**
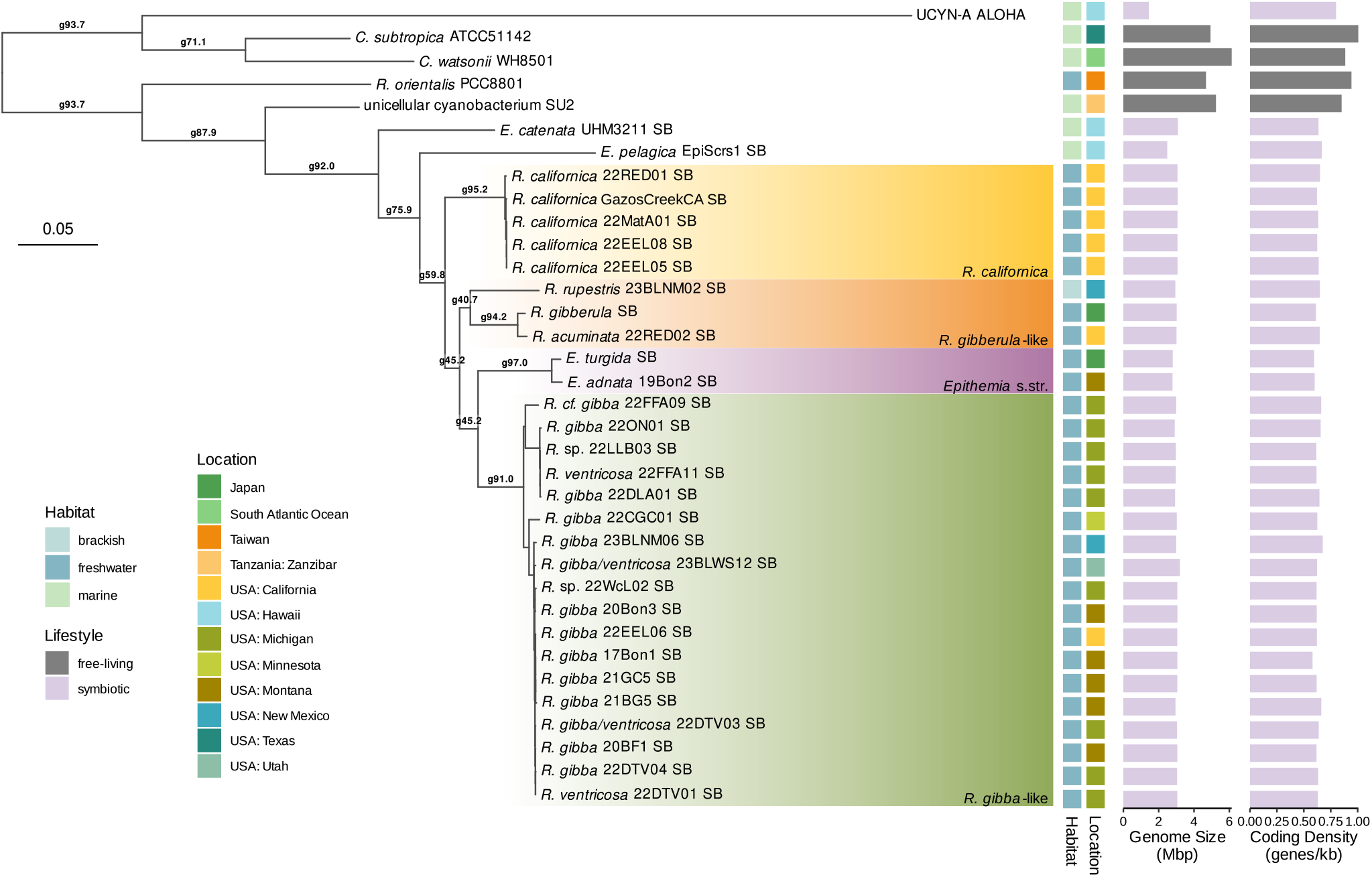
A maximum likelihood, genome-wide phylogeny of SBs and five outgroups (four free-living cyanobacteria, and cyanobacterial endosymbiont UCYN-A ALOHA). The phylogeny was constructed from a concatenated alignment of 398 amino acid sequences using the Q.plant+F+I+G4 model of evolution and 1000 bootstrap replicates. Habitat, lifestyle, genome size, and coding density are indicated by tiles and bars according to the legend. Gene concordance factor (gCF) values for major nodes of interest leading to the different SB clades and outside of the SB group are shown as numbers along branches (formatted gXX.X). All gCF and gene discordance factor values for all branches are in Supplemental Figure 2. All bootstrap values = 100 except for: branch leading to 22EEL05 and 22EEL08 = 99.7, branch leading to 22DTV01 and 22DTV04 = 98.1, branch leading to 21BG5 and 21GC5 = 97.6, branch leading to 22DLA01 and 22FFA11 = 99.5.

SB genome size ranged from 2.4 Mega base pairs (Mbp) to 3.2 Mbp, with a GC content of 33.4% to 35.3% (Figure 1, Supplemental Table 1). SB chromosomes have 1,656 to 2,021 (mean = 1,889, median = 1,909) annotated protein coding sequences (CDS) and an estimated coding density of 0.58 to 0.68 genes per kilobase (mean and median = 0.63 genes per kilobase; Figure 1), which is much lower than typical coding density for free-living bacteria (Mira et al., 2001; Giovannoni et al., 2005). This low coding density can be attributed largely to intergenic regions with degraded gene fragments that no longer have intact open reading frames for annotation (Supplemental Table 1). These results agree with models of endosymbiont genome evolution that predict that, during early stages following host restriction, coding density reaches a minimum and number of pseudogenes peak (McCutcheon and Moran, 2012; Lo et al., 2016).

All closed SB genomes have two copies of the *rrn* operon (16S, 23S, and 5S rRNA genes). We expect this holds true for fourteen of the draft assemblies where the contig encoding the *rrn* operon and tRNA-Ile assembled into a single contig (the “*rrn* contig”) with double the coverage of other SB contigs. However, we found evidence for greater than two copies in five draft assemblies based on 3-6× elevated *rrn* contig coverage compared with other contigs. There are also five draft assemblies where the *rrn* contig has the same coverage as the rest of the assembly, which suggests only a single copy of the *rrn* contig in these genomes. The apparent variation in *rrn* contig copy number did not carry a strong phylogenetic signal, but were distributed across the SB phylogeny (Supplemental Table 1). Overall, this suggests more variation in *rrn* copy number than previously recognized.

A small plasmid has also been reported in several SB genomes (Nakayama and Inagaki, 2017; Abresch et al., 2024). We identified this plasmid as a distinct contig in all metagenome assemblies. The *∼*6,000 bp plasmid contains five genes, including ferrous (Fe^2+^) iron import proteins, *feoAB*, and aquaporin Z, *aqpZ*, except for the *Epithemia catenata* SB plasmid, which is approximately three times longer than all other plasmids and encodes 11 genes. Based on relative coverage, we estimate there are 1-2 copies of the plasmid per SB chromosome (Supplemental Table 1). Two previously published assemblies, *E. pelagica* EpiScrs1 SB (GenBank RefSeq: GCF 947331815.1) and *R. californica* GazosCreekCA SB (published as *E. clementina* SB in Moulin et al. (2024)), do not report the presence or absence of this plasmid, but given our result we suspect it is present in all SBs.

Models of endosymbiont genome evolution predict that proliferation of mobile elements drives genome reduction and leads to frequent chromosome rearrangements and low levels of synteny in recently host-restricted endosymbionts (McCutcheon and Moran, 2012). In contrast to this model, we found that SB genomes have no intact transposases and chromosomes are highly syntenic (Supplemental Table 2, Supplemental Figure 1), consistent with previous studies (Nakayama and Inagaki, 2017; Abresch et al., 2024). Genomes from closely related free-living cyanobacteria have many transposases and structural rearrangements are common (Bandyopadhyay et al., 2011; Xu et al., 2016); therefore, SBs likely lost all transposases early in the genome reduction process. Whether the early loss of transposases was neutral or selectively favored is unknown. The early loss of all mobile elements is unusual in younger endosymbionts. The loss of transposases combined with high levels of synteny in SB genomes supports the hypothesis that proliferating mobile elements drive genome instability during the early stages of endosymbiont genome evolution seen in other systems (McCutcheon and Moran, 2012).

### Genome-wide phylogeny for spheroid bodies

We reconstructed a genome-wide maximum likelihood phylogeny for SBs and five outgroup taxa (four closely-related, free-living cyanobacteria and UCYN-A ALOHA) using 398 single-copy orthologs (Figure 1). This phylogeny has high bootstrap support at all nodes. SBs formed a clade, which supports a single origin of the SB endosymbiosis. With strong support, the phylogeny identifies the closest, free-living relative as “unicellular cyanobacterium SU2” (hereafter, SU2), a marine cyanobacterium from the coast of Zanzibar, Tanzania. The closest well-described relative is *Rippkaea orientalis* PCC 8801, a freshwater unicellular cyanobacterium. This agrees with our previously published genome-wide tree using only four SBs (Abresch et al., 2024). By contrast, single- and two-gene phylogenies (using only the 16S rRNA gene, *nifH*, or both) identify either *Crocosphaera subtropica* ATCC 51142 or the UCYN-A lineage as the sister to SBs, but lack strong support at relevant internal nodes (Prechtl et al., 2004; Schvarcz et al., 2022; Moulin et al., 2024). Our placement of SBs within *Rippkaea* instead of *Crocosphaera* supports the independent evolution of SBs and the UCYN-A group (Figure 1).

SBs resolved into four major clades which largely agree with the topology of published host phylogenies (Ruck et al., 2016; Kociolek et al., 2024). As such, we have labelled these clades based on their host groups (*R. californica* group, *R. gibberula*-like *sensu* Kramer 1988, *Epithemia sensu stricto* [s.str.] group, and *R. gibba*-like group; Figure 1). This suggests predominantly, if not exclusively, vertical transmission of the SB during host reproduction. Although the broader topology between hosts and SBs matches, a better understanding of host evolutionary history is needed to clarify concordance between hosts and SBs within clades.

The phylogeny also shows a single transition from marine to freshwater habitats (Figure 1). SU2 and the earliest branching SBs are from marine habitats, and later branching SBs are from freshwater or inland brackish habitats, suggesting that diatom hosts and SBs transitioned together to freshwater ecosystems and subsequently diversified. Only two SBs from marine host species (*E. catenata* UHM 3211 SB, this study; and *E. pelagica* EpiScrs1 SB, (Schvarcz et al., 2022)) have been described. More exploration of SBs and hosts from marine habitats will be informative in understanding the evolution of Rhopalodiaceae and their transition from marine to freshwater environments.

Gene trees also generally support the species tree as shown by gene concordance factor (gCF) values. Longer branches have high gCF values (Figure 1). As expected (Minh et al., 2020), gCF values are much lower in very short branches, such as within the *R. gibba*-like or *R. californica* groups (Supplemental Figure 2). However, gCF values are slightly low at branches where major SB clades diverge (Figure 1). We found much of this discordance is due to varied placement of either an entire clade of SBs (e.g. the *R. gibberula*-like clade appears before the *R. californica* group), or specifically movement of 23BLNM02 SB, which is within the *R. gibberula*-like group in the species tree (Supplemental Figure 3). Gene tree discordance could arise from biological causes such as incomplete lineage sorting (ILS) of ancestral polymorphism, e.g., following rapid speciation events, which we believe contributes to discordance seen in the SB phylogeny. Alternatively, gene tree discordance could come from low phylogenetic signal or sequence saturation that overshadows true phylogenetic signal (the latter is unlikely for short branches). Because the gDF1 and gDF2 values across these branches are low and comparable, discordance is not due to a systemic bias towards a particular alternative topology (Supplemental Figure 2), which would result from horizontal gene transfer (HGT) event(s) between more distantly related SBs.

### Sampled SB genomes provide a robust estimate of the core genome

To better understand the genome reduction process and to identify conserved and potentially idiosyncratic metabolic capabilities among SBs, we estimated SB core and pan genomes. We focused our analysis on SB assemblies with fewer than 50 contigs (N = 22 genomes; see Materials and Methods) to minimize the erroneous exclusion of genes from the core. After estimating the pangenome from 22 SBs, we used rarefaction to estimate how many genes would be lost from the core (i.e. the size of the “true” core) and how many more new genes we would likely find (e.g. whether the pangenome is open or closed) with additional SB genomes.

Overall, we found that most genes are either in all 22 genomes or fewer than 3 genomes (Figure 2A, Supplemental Table 3). There were 3,164 gene clusters in the pangenome. The core genome contains 1,285 genes; therefore *∼*75% of genes in any one SB genome are core genes (Supplemental Figure 4). Of the remaining pangenome, 60% (N = 1,119) were in 3 or fewer genomes, and 670 genes were in anywhere from three to 20 genomes (Figure 2A).

**Fig. 2:**
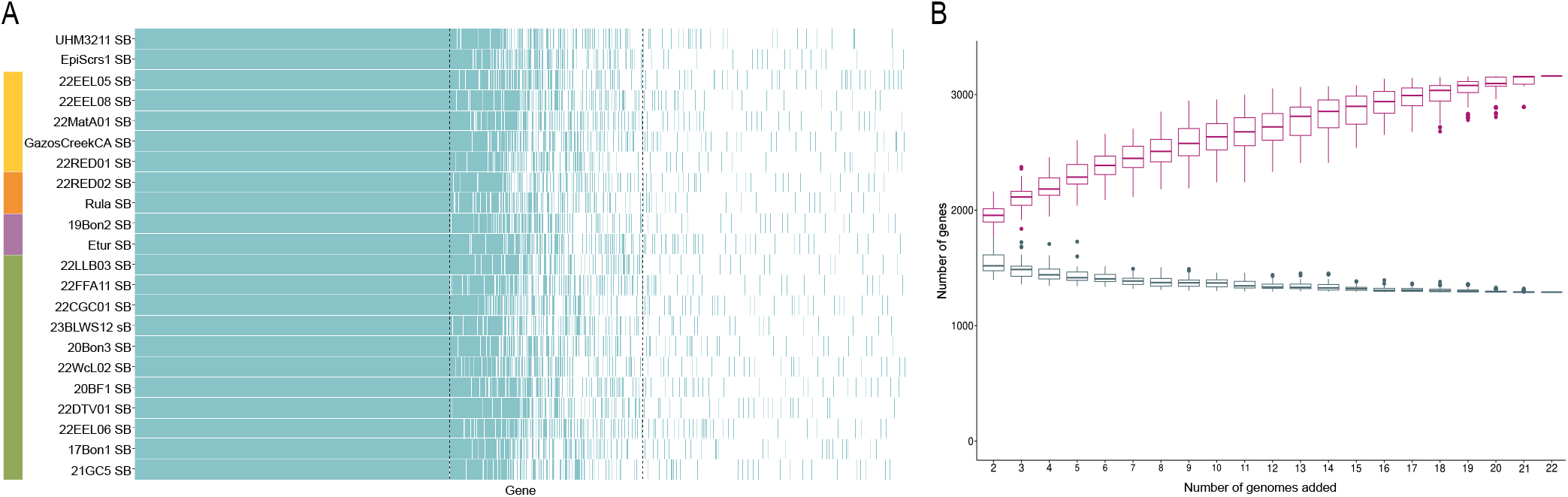
A) Gene presence and absence in 22 SB genomes. Colored bars on the left correspond= to highlighted SB clades in Figure 1. B) Rarefaction of SB core (lower; blue) and pan genomes (upper; pink) based on 22 genomes and 100 permutations.

A binomial mixture model estimated the true SB core genome to be 1,230 genes, only 55 genes fewer than the core estimated from 22 genomes (Figure 2B). This suggests that the estimated core is an accurate estimate of the “true” SB core genome, toward which individual SB genomes are converging. That is, including additional SB genomes would not result in a significantly smaller core genome size. Theory predicts that the size of an endosymbiont’s core genome is a product of both its metabolic functions and the ability of selection to prevent deleterious gene loss (Bennett et al., 2024). Supportive of this, the number of core genes in SBs is similar to the number of core genes found in other N-fixing cyanobacterial endosymbiont lineages; for example, the UCYN-A core genome contains an estimated 1,070 genes (Kantor et al., 2024).

In contrast, the model estimated the SB pangenome to be 4,989 genes, over 1,800 more than the number of genes found in only 22 genomes (Figure 2B). Generally, host-restricted and vertically transmitted endosymbionts do not gain any new genes via HGT (McCutcheon, 2021) and we do not see any evidence of this in SBs. Therefore, early endosymbiont pangenomes reflect the ancestral gene content. Indeed, the estimated size of the SB pangenome is roughly the same size as the number of genes in free-living members within Aphanothecaceae (typically *∼*5,000 genes), although it is slightly larger than the closest free-living relatives sampled (*R. orientalis* PCC 8801 = 4,409 protein coding genes; SU2 = 4,460 protein coding genes; Supplemental Table 1).

We expect that, in late-stage endosymbionts, the size of the pangenome will approach that of the core genome, as non-essential genes are lost over time, whereas early endosymbionts are still losing genes stochastically and the pangenome will be much larger than the core. Therefore, a smaller pangenome is reflective of an endosymbiont being further along in the genome reduction process, even in endosymbionts with similar core genome sizes. While SBs and UCYN-A lineages have similar core genome sizes, the UCYN-A pangenome contains only approximately 1,400 genes (Kantor et al., 2024)—much smaller than the SB pangenome. From this, we conclude that UCYN-A is much further along in the genome reduction process than the SB.

### Spheroid body core metabolic capacity

Genes retained across all endosymbiont genomes reflect their conserved metabolic functions (McCutcheon, 2021). Conversely, entire metabolic pathways which are only retained within a subset of SB lineages could indicate divergence of SB function among hosts. The presence or absence of genes typically lost from endosymbiont genomes over long-term host restriction could also inform the extent to which SBs are integrated with their host metabolism. Here, we focus primarily on the metabolic capacity of the core genome, rather than the capacity of individual SBs.

We examined SB core metabolic capacity by assigning KO (Kyoto Encyclopedia of Genes and Genomes [KEGG] Orthology) IDs to core genes. To differentiate between ancestrally incomplete pathways versus those lost during SB evolution, we compared gene content of the SB core to both *R. orientalis* PCC 8801 and SU2. Of the 1,285 core SB genes, 850 (66.5%) were assigned at least one KO ID. All but four of these genes best matched an ortholog within Cyanobacteria and 1,069 of these best matched *R. orientalis* PCC 8801 specifically, which agrees with our phylogeny findings. Generally, we found that SBs share the same complete pathways or are missing various parts of metabolic pathways in a manner that suggests stochastic degradation. For brevity, we highlight a few key pathways here, but we provide an extended overview of metabolic pathways in Supplemental Figure 5.

All essential N-fixation genes have been retained. *nifJ*, which is used in N-fixation under iron limitation, has been lost from SBs and other N-fixing cyanobacterial endosymbionts (Kantor et al., 2024; Grujcic et al., 2025), likely reflecting a more stable intracellular environment (Grujcic et al., 2025). Amino acid biosynthetic pathways are comparable to free-living relatives (Supplemental Figure 5). As expected for endosymbionts, SBs have also lost the genes for many transporters, including iron (Fe) (III) and nitrate uptake, but they retain genes for Fe (II) and molybdate uptake, both required for N-fixation.

Many core carbon metabolic pathways are incomplete (Supplemental Figure 5). Glycolysis is missing phosphofructokinase (*pfk* ), and the citric acid cycle is missing succinate dehydrogenase and malate dehydrogenase. The oxidative pentose phosphate pathway remains complete, while the non-oxidative phase is incomplete. It is possible that carbon is prioritized to move through the oxidative pentose phosphate pathway instead of the citric acid cycle to promote NADPH regeneration, which is required for N-fixation (Bothe et al., 2010).

The SB core is also missing pathways for synthesizing many vitamins and cofactors. Both chlorophyll and phylloquinone synthetic pathways are completely absent. SBs have also lost the genes for molybdenum (Mo) cofactor synthesis. This likely allows all Mo available to be used in Fe-Mo cofactor synthesis, which is required for N-fixation. As previously noted (Nakayama and Inagaki, 2017; Abresch et al., 2024; Chang et al., 2025), the only major difference in metabolic pathways among SB genomes is in the presence or absence of the vitamin B_12_ (pseudocobalamin) biosynthetic pathway, which is complete in most SBs, but almost completely lost in SBs from *Epithemia s. str*. spp., suggesting divergence in SB function across hosts.

### Universally absent “essential” genes are potential points of host control of SB growth

Endosymbionts often partially or entirely lose genes required for the essential processes of DNA replication and repair and for cell wall (peptidoglycan; PG) synthesis (Bennett and Moran, 2015; McCutcheon, 2021). This recurring pattern of loss suggests that DNA replication and PG synthesis and degradation may be common loci for hosts to establish control over endosymbiont growth and division. Additionally, products of PG degradation are often immunogenic and, by limiting the release of these molecules, endosymbionts could prevent a negative reaction from the host (Otten et al., 2018). Contrary to this, we found that both PG synthesis and DNA replication and repair remain largely intact in SBs, with two major exceptions: *dnaA* and *mltA* (Supplemental Figure 5). Despite these genes typically being considered essential for bacterial cell growth (Heidrich et al., 2002; Kaguni, 2006; Lamers et al., 2015), both are universally absent in SBs. These genes are therefore promising candidates for host control over SB growth and division.

*dnaA* is typically considered essential for initiating genome replication (Kaguni, 2006), although *dnaA* dependency may be less strict in cyanobacterial lineages (Ohbayashi et al., 2020). This gene has been reported missing in diverse bacterial endosymbionts, including other cyanobacterial endosymbionts (Muñoz-Marín et al., 2019), the gamma-proteobacterium *Wigglesworthia* (Akman et al., 2002), and the bacteriodete *Blattabacterium* (Whittle et al., 2021). The repeated loss of *dnaA* makes it particularly interesting as a commonly evolved mechanism for hosts to gain control over endosymbiont division, possibly using host encoded proteins for initiation of replication and allowing hosts to control the costs of symbionts (e.g. number of symbionts per host cell, metabolic cost, etc.) (Klasson and Andersson, 2006). The lytic transglycosylase MltA is involved in PG degradation and is a key protein in maintaining cell wall integrity in bacteria (Heidrich et al., 2002; Lamers et al., 2015; Smith et al., 2022). In many studies, the deletion of *mltA* adversely impacts cell division, growth, and both membrane architecture and permeability (Adu-Bobie et al., 2004). In an endosymbiotic bacterium infecting silkworm moth ovary cells, the deletion of *mltA* limited intracellular growth and cytotoxicity (Nakamura et al., 2021). The loss of *mltA* in SBs could have similarly limited growth within the host cell and reduced the costs to the host associated with supporting SBs, allowing SBs to establish a stable, lasting presence within hosts.

### Strength of selection on core genes

We next examined how selection is acting on core genes. Core genes contribute to essential SB functions and are thus expected to be under relatively strong purifying selection, provided that selection strength outweighs the strength of genetic drift. Generally, drift is stronger in endosymbionts due to low effective population size driven by population bottlenecks during host cell division (McCutcheon and Moran, 2012). Whether or how the strength of selection changes over time on core genes in early endosymbionts versus late-stage endosymbionts is not well understood. The strength of selection versus drift may also vary in endosymbionts from single-celled hosts versus multicellular hosts; multicellular hosts generally harbor more endosymbionts per host and can pass more endosymbionts on during host reproduction, which could affect the ability of selection to maintain protein function (Husnik et al., 2021). A higher number of endosymbionts may allow for stronger selection within a host and more endosymbionts passed on during cell division would change the dynamics of the population bottleneck. Whereas an endosymbiont within a single-celled host may have fewer endosymbionts, which are subjected to bottlenecks every cell division, though selection strength and fitness of endosymbionts may be more directly tied to the host’s in this scenario. To evaluate the strength of selection on core SB genes, we estimated rates of non-synonymous substitutions per non-synonymous site to synonymous substitutions per synonymous site (dN/dS) across 11 SB genomes representing SB diversity, where genes under purifying selection will have a dN/dS less than one.

We found that there is moderately strong purifying selection on core genes tree-wide with a mean dN/dS of 0.173 and median of 0.166 (Figure 3, Supplemental Figure 6). While the average dN/dS is relatively low, the range of values is larger than those observed in older endosymbionts with more reduced genomes (Silva and Santos-Garcia, 2015; Boscaro et al., 2022; Siozios et al., 2024). Overall, our results suggest that although the strength of genetic drift is generally higher in endosymbionts, selection to maintain a functional core is strong.

**Fig. 3:**
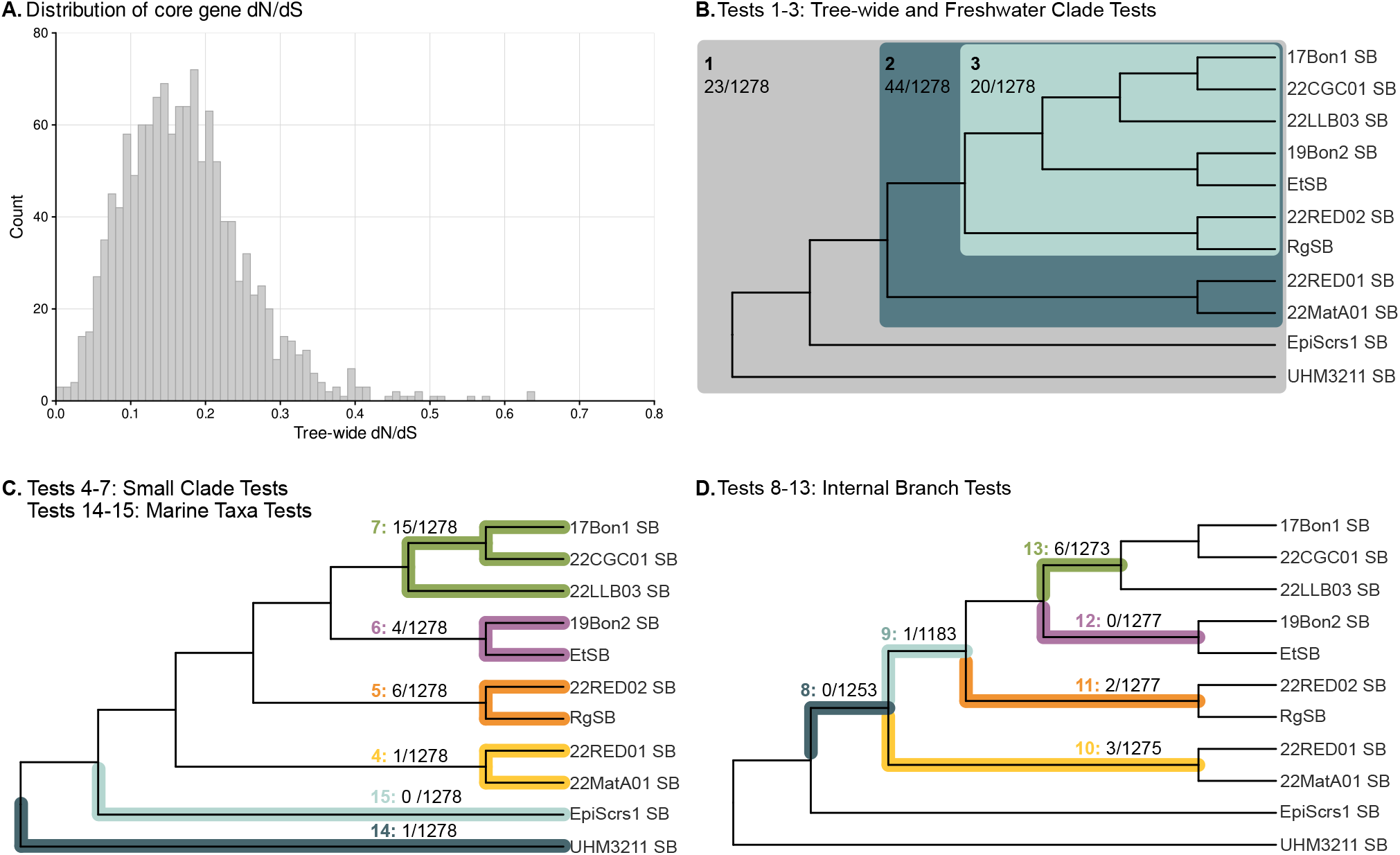
A) Histogram of estimated dN/dS for all SB core genes. B), C) and D) Likelihood ratio tests used to test for positive selection in SB core genes. Branches tested for positive selection in each test are highlighted. The number of positive results after false discovery rate correction out of total number of tests for each test are shown. All tests were run using an unrooted phylogeny.

Conversely, genes that are directly involved in host interactions are expected to be under positive selection (Bennett and Moran, 2015). We tested for episodic positive selection to identify candidate genes in two ways: 1) tree-wide, and 2) along internal branches and within the different SB clades (Figure 3). We took a likelihood approach using branch-site models (HyPhy BUSTED) to identify genes with evidence that at least one site has experienced positive selection during diversification along the tested branch(es). We ran a total of 15 tests: a tree-wide test (t1), and 14 tests which designated different branch(es) of the phylogeny as test (foreground) branches. Tests t2 and t3 tested for positive selection within SBs from freshwater hosts, including or excluding the earliest branching freshwater clade (the *R. californica* group), respectively. Finally, we tested within SB clades (t4 through t7), internal branch tests (t8 through t13), and marine taxa tests (t14 and t15), as shown in Figure 3 and Supplemental Figure 6.

Across all tests, we identified a total of 54 genes (across 126 significant results) with evidence of episodic positive selection. The tree-wide test found evidence of episodic positive selection for 1.8% of genes (N = 23, Figure 3). All genes that were found to be significant in the tree-wide test were also significant in at least one other test, except one (*glgB* ). Furthermore, all but eight genes were identified in at least one of t1, t2, or t3 (Supplemental Table 4). This suggests that, for the majority of genes, a tree-wide signal for positive selection is the product of bouts of selection along specific branches rather than continuous selection during SB diversification (i.e., as would be expected for an ongoing evolutionary arms race).

Except for t7, all tests of individual internal branches or individual clades yielded six or fewer significant results, including three tests that yielded no significant results (Figure 3, Supplemental Table 4). Test 7, which tested for genes under positive selection within the *R. gibba*-like clade, identified 15 significant genes. Interestingly, *R. gibba*-like hosts also have a strict number and positioning of SBs (Geitler, 1977), suggesting a higher level of integration with SBs compared to other host species. Our selection results indicate there may be more direct SB-host interactions in the *R. gibba*-like clade of SBs compared to the other SB clades.

### Core genes under positive selection are possible points of direct host-SB interaction

Next, we examined the functionality of the putatively positively selected genes. In recently restricted endosymbionts, the endosymbiont and host genes that directly interact with each other are predicted to evolve under arms race dynamics, which can be seen as positive selection (Bennett and Moran, 2015; Chong et al., 2019; McCutcheon, 2021). Thus, we expected to find genes encoding peripheral proteins, including transporters and genes involved in making cell membrane/cell wall components, to be highly represented, as they are likely important to the symbiosis and directly involved in host-SB interactions in a physical sense.

Of the 54 genes significant for positive selection, 52 had a closely related free-living homolog to compare to in relevant databases. For 44 of these, the closest related sequence was from *R. orientalis* PCC 8801 (SU2 is not available in these databases for comparison). 49/54 genes had an identifiable conserved domain, 38 were assigned a KO ID, and 29 genes had a predicted protein subcellular location (Supplemental Table 5).

In general agreement with theory, one third of these genes (N = 21) are membrane-associated or involved in membrane and cell wall metabolism: seven predicted transporters, three genes involved in signal transduction, ten genes involved in membrane synthesis or PG metabolism, and one gene putatively involved in regulating cell division. Eleven genes significant for positive selection are related to genetic information processing: eight involved in translation, two involved in transcription, and one involved in DNA repair and replication. While we did not explicitly expect genes involved in genetic information processing to be highly present in our results, genes in these categories have also been found to be under positive selection in the well-studied aphid endosymbiont *Buchnera*, which has a highly reduced genome, although the exact genes are not the same (Chong et al., 2019).

The most surprising result was that three *nif* genes were significant for positive selection: two core nitrogenase genes (*nifH* and *nifK* ) and *nifS*, which is involved in iron-sulfur cluster synthesis for N-fixation. Since these genes contribute to the core function of SBs, we expected them to be understrong purifying selection to maintain functionality. One possibility is that SBs are adapting to their new intracellular environment. For instance, SBs may experience increased oxidative stress. Nitrogenase is irreversibly inactivated by oxygen, these two processes are typically temporally separated in unicellular cyanobacteria (Bothe et al., 2010). However in SBs, N-fixation occurs concurrently with host photosynthesis (Schvarcz et al., 2022; Abresch et al., 2024). Consequently, there may have been the potential for selection to favor a more oxygen-insensitive nitrogenase.

## Conclusion

The current model of endosymbiont genome evolution predicts that endosymbiont genomes are initially unstable and lose genes rapidly due to deletions resulting from recombination between proliferating mobile elements, the “active” genome reduction phase. Eventually, genomes lose all mobile elements and become more stable, i.e., the “late” genome reduction phase (McCutcheon, 2021). In contrast to this model, we found that SBs have many predicted pseudogenes and very low coding density. Yet they have no transposases and are relatively stable structurally. The current model would predict that SBs would either have significantly more mobile elements or much smaller genomes. Additionally, the SB core genome is small and well-predicted by the current sampling of SB genomes, while the SB pangenome is large and not well represented by the current sampling. Model estimates suggest the “true” SB pangenome likely still reflects ancestral genome content, as opposed to the pangenome representing only a subset of the ancestral genome. The SB core metabolic capacity also agrees with expectations based on other endosymbionts. Based on the data presented here, we suggest SBs are still in the “active” genome reduction phase, with stochastic losses still occurring across SB lineages, and the stability of SB genomes is instead due to the early loss of all transposases.

The SB genome-wide phylogeny generally reflects host species relationships with no obvious history of HGT between SBs or transfer of SBs between distantly related hosts. A better understanding of host evolutionary history is needed to understand within-host clade dynamics of SB transmission, e.g. generation of novel host-SB genotype combinations through varied uniparental inheritance during sexual reproduction, as explored in Kamakura et al. (2021).

How SBs and hosts interact, or how much control hosts have over SBs, is yet unknown. We identified putative points of host control and SB-host direct interactions in two ways. Here, we identified 2 key gene absences that may be points of host control over SB division and 54 core SB genes undergoing positive selection, many of which encode peripheral proteins or genes involved in membrane or cell wall synthesis. This greatly expanded SB data provided here will guide further investigations of how the diatom hosts and SBs are genetically and metabolically integrated. Overall, this study furthers understanding the key processes in the early stages of endosymbiont evolution and expands our knowledge of endosymbionts in unicellular hosts which will allow comparisons of endosymbiont evolution across diverse host types.

## Materials and Methods

### Cell isolation and culturing

The 22 new SB-containing diatom strains presented in this paper were isolated from environmental samples collected between August 2020 and May 2023. Information on sampling locations and dates are available in Supplemental Table 1. Strains were isolated and maintained in N-free media as described in Abresch et al. (2024). Cultures are unialgal and uni-eukaryotic, but not axenic.

### DNA extraction and sequencing

Cells were pelleted and resuspended in 900 *µ*L TE and 100 *µ*L of 1% Triton-X (in TE) to wash away exogenous bacteria before DNA extraction. DNA was extracted using the Qiagen DNeasy PowerBiofilm kit according to manufacturer instructions. Metagenomic libraries were prepared using the Illumina DNA Prep tagmentation kit. Paired-end, 150 bp short read sequencing was performed on the NextSeq2000 platform (Illumina). Library preparation and sequencing was performed by SeqCoast (Portsmouth, NH, USA).

### Genome assembly and annotation

We generated *de novo* metagenome assemblies using SPAdes (Bankevich et al., 2012) version v3.12.0 or v3.15.5 with the `--meta` option. Contigs shorter than 1,000 bp or *<*1× coverage were removed from the metagenomic assembly. SB sequences were identified by sequence similarity to previously published SB genomes (chromosome and plasmid) using BLAST v2.15.0+ (Camacho et al., 2009) and extracted from the metagenome assembly. Identified SB scaffolds were compared to the full core nucleotide database on NCBI and confirmed to have the closest identity to another SB genome. SB assemblies with *<*100 contigs were manually inspected by aligning raw reads to the full metagenomic assemblies in Bowtie v2.5.1 (Langmead and Salzberg, 2012) and visualizing in Integrative Genome Viewer (Robinson et al., 2011). SB assemblies in *>*100 contigs were considered draft assemblies and not further manually inspected.

All SB genomes were annotated by PROKKA v1.14.6 (Seemann, 2014) using default settings with `--compliant` and `--rfam` to detect non-coding RNAs. Because PROKKA does not identify pseudogenes, we then refined SB chromosomes using pseudofinder v1.10 (Syberg-Olsen et al., 2022) to identify putative pseudogenes using commands `-hc 100 -dev -nbl` compared to a custom BLAST+ database comprised of 17 free-living cyanobacterial genomes from the order Chroococcales available on the NCBI RefSeq database. Previously published genomes (EturSB, RulaSB, EclemSB, EpiScrs1SB, Rgib 17Bon1 SB, and Eadn 19Bon2 SB) were reannotated following these methods for consistency in comparative analyses. Additionally, genome completeness was assessed using CheckM (Parks et al., 2015) for the core gene set from marker lineage Chroococcales. For SB draft assemblies, ribosomal copy number and tRNA count was estimated based on coverage of the contig containing the ribosomal operon compared to coverage the other SB chromosome contigs in the assembly. In all instances, the *rrn* contig only encoded the ribosomal subunits and the tRNA that is encoded between the SSU and LSU.

Gene function prediction and metabolic pathway reconstruction were done using eggNOG v.emapper-2.1.9 (Huerta-Cepas et al., 2019) using the DIAMOND database to assign KO (Kyoto Encyclopedia of Genes and Genomes [KEGG] Orthology) IDs to annotated SB genomes using default settings. All but eleven core SB genes met the minimum EggNOG search criteria.

### Phylogenies

We used Orthofinder (Emms and Kelly, 2019) to identify, align, and concatenate 398 single-copy ortholog protein sequences (113,427 amino acid sites) shared by all 30 SB taxa and 5 outgroup taxa. A maximum likelihood phylogeny with 1,000 UltraFast bootstrap replicates was constructed in IQTree2 v2.2.2.6 (Nguyen et al., 2015) using the Q.plant+F+I+G4 model of evolution selected by ModelFinder (Kalyaanamoorthy et al., 2017). Tree branches were tested by SH-like aLRT with 1000 replicates (Guindon et al., 2010). We also constructed gene trees of all 398 single-copy orthologs and used these to generate gCF values in IQtree2.

### Core and pangenome estimation

We used the 22 best SB assemblies to estimate the SB core and pan genome in Roary v3.11.2 (Page et al., 2015) with a minimum BLASTp identity of 75% (-i 75). We then manually examined core genes in gene groups that had 1) *>*200 amino acid difference between the minimum and maximum protein length, or 2) a minimum protein length *<*80% of maximum protein length. Additionally, we manually examined the initial output to ensure genes missing from only one or two SBs were true absences and not due to partial genes located at contig breaks.

The majority of the manually inspected gene groups contained only one or two SBs whose ortholog was much shorter than all other SB copies. To be conservative in our downstream analyses, we minimized erroneous exclusion of genes from the core during manual inspection to not underestimate SB metabolic capacity (i.e. manual changes were conservative in favor of gene presence rather than absence). In instances where one gene was shorter than the rest in the group, the “short” gene was considered intact if the gene was found next to a contig break in the correct direction and otherwise present in all other SBs. Due to manual changes made, we have provided the final table of gene presence and absence used here in Supplemental Table 3. Rarefaction analyses and core and pan genomes were estimated using the micropan package and custom R code in R version 4.5.1 (see Data Availability Statement).

### Selection analyses

We estimated the strength of selection and tested for episodic positive selection with HyPhy BUSTED (Branch-site Unrestricted Statistical Test for Episodic Diversification) v2.5.74(MP) (Murrell et al., 2015), which fits gene alignments to two different codon models along a phylogeny: a constrained model, without positive selection (i.e. maximum change rate limited to *≤* 1); and an unconstrained model, where positive selection is allowed but not required. If the unconstrained model fits the data significantly better and at least some codon sites are significantly assigned to the positive rate class, then there is evidence that at least one site has experienced positive selection during diversification along the tested branch(es). We ran 15 tests of positive selection on 1,278 core genes. Overall tree-wide dN/dS estimates were calculated in the tree-wide test (t1) which also tested whether positive selection appeared at least somewhere within the tree. Tests two through fifteen (t2 - t15) specified a subset clade or internal branch as foreground within the model using the `--branches` option. For t2 – t15, genes where the foreground branch length(s) were zero were not tested. We accounted for the possibility for multi-nucleotide mutations, which can lead to false positives (Venkat et al., 2018), using FitMultiModel.bf built within HyPhy. For the 76 genes with possible multi-nucleotide mutations, we reran these through HyPhy BUSTED with the option `--multiple-hits “Double+Triple”`. Finally, for each test, we filtered out genes for which the models did not converge (likelihood ratio test *≤* 0) and corrected the remaining p-values for multiple tests using Benjamini-Hochberg correction with a false discovery rate of 0.1 using the R package qvalue (Storey et al., 2025). For further classification of genes significant for episodic positive selection, we used a combination of conserved domains within SB protein sequences identified by blastp online and KEGG and UniProt database information for highly similar homologs (e.g., in *R. orientalis* PCC 8801).

## Data Availability Statement

For BioRxiv preprint, data availability TBD. All BioSamples and genome data will be made publicly available via NCBI under the BioProject Accession Number PRJNA1379625. GenBank accession numbers for SB assemblies are listed in Supplemental Table 1. Custom R code and additional data files are available on

GitHub at https://github.com/cyanochic/SBComparativeGenomics_public.

## Acknowledgments

Computational resources and support from the University of Montana’s Hellgate Research Cluster contributed to this research. The Hellgate Research Cluster’s role in this publication was provided free of charge or obligation. *E. catenata* UHM3211 cells were provided by Grieg Steward at University of Hawaii Manoa. Original sample material for *R. gibba* 22CGC01 was provided by Siri Davidson.

