## Supplemental Material for "Diatom Endosymbionts have Shrinking but Stable Genomes Despite Low Coding Density"

### Supplemental Tables

**Supplemental Table 1:** Summary of genome content for SBs and close, free-living relatives: *Rippkaea orientalis* PCC 8801 and unicellular cyanobacterium SU2. We additionally provide information on the alternate genome annotation method PGAP, alongside the PROKKA plus pseudofinder annotation method we report on in the main text. * = number of rRNA gene clusters estimated based on relative coverage of the *rrn* contig; ** = only based on assembly annotation, no coverage information included.

**Supplemental Table 2:** Best transposase hits for SB genomes from ISFinder.

**Supplemental Table 3:** Gene presence and absence table after manual curation from Roary base output. Cells filled with a dash ("-") indicate manual confirmation of gene absence. The `locus_tag` IDs correspond to `locus_tag` IDs found in the file `allSB_prokka.faa` provided in the GitHub repository for the main paper (See Data Availability Statement).

**Supplemental Table 4:** Selection results for all genes in SB core, note that the p-value included is the raw p-value and does not account for FDR correction.

**Supplemental Table 5:** Functional summary of genes significant for positive selection in at least one test.

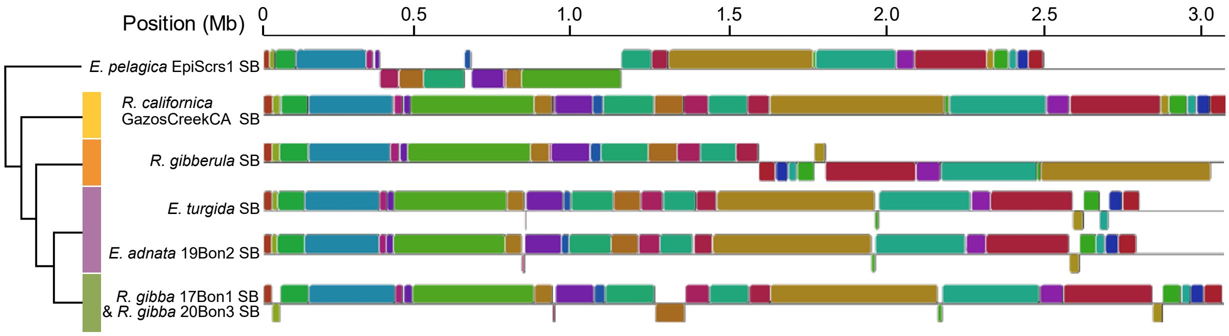


Figure S1: SB chromosome synteny of closed SB assemblies constructed with progressiveMauve after SB chromosomes were manually adjusted to start at the same ribosomal operon. Both *R. gibba* genomes have been collapsed into one line as they did not have any syntenic differences. Colored bars on the cladogram match those in main text figures.

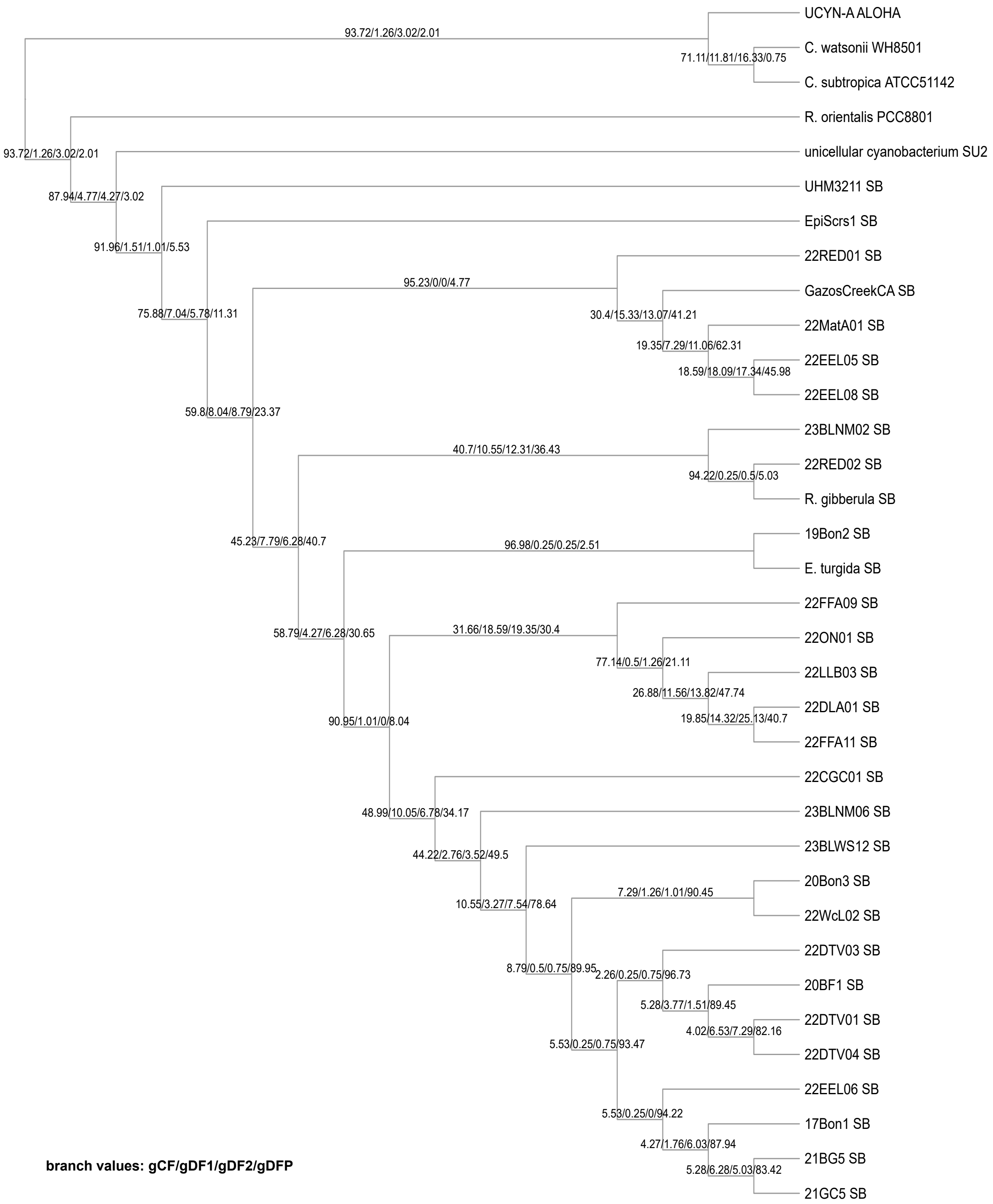


Figure S2: SB phylogeny without branch lengths to scale showing gene concordance factor (gCF) and all gene discordance factor (gDF1, gDF2, and gDFP) values formatted as gCF/gDF1/gDF2/gDFP. gDF1 and gDF2 represent the two alternate topologies and gDFP is from paraphyly. See Minh et al., 2020 (referenced in main text) Figure 1 and related text for further description of gCF and gDF values.

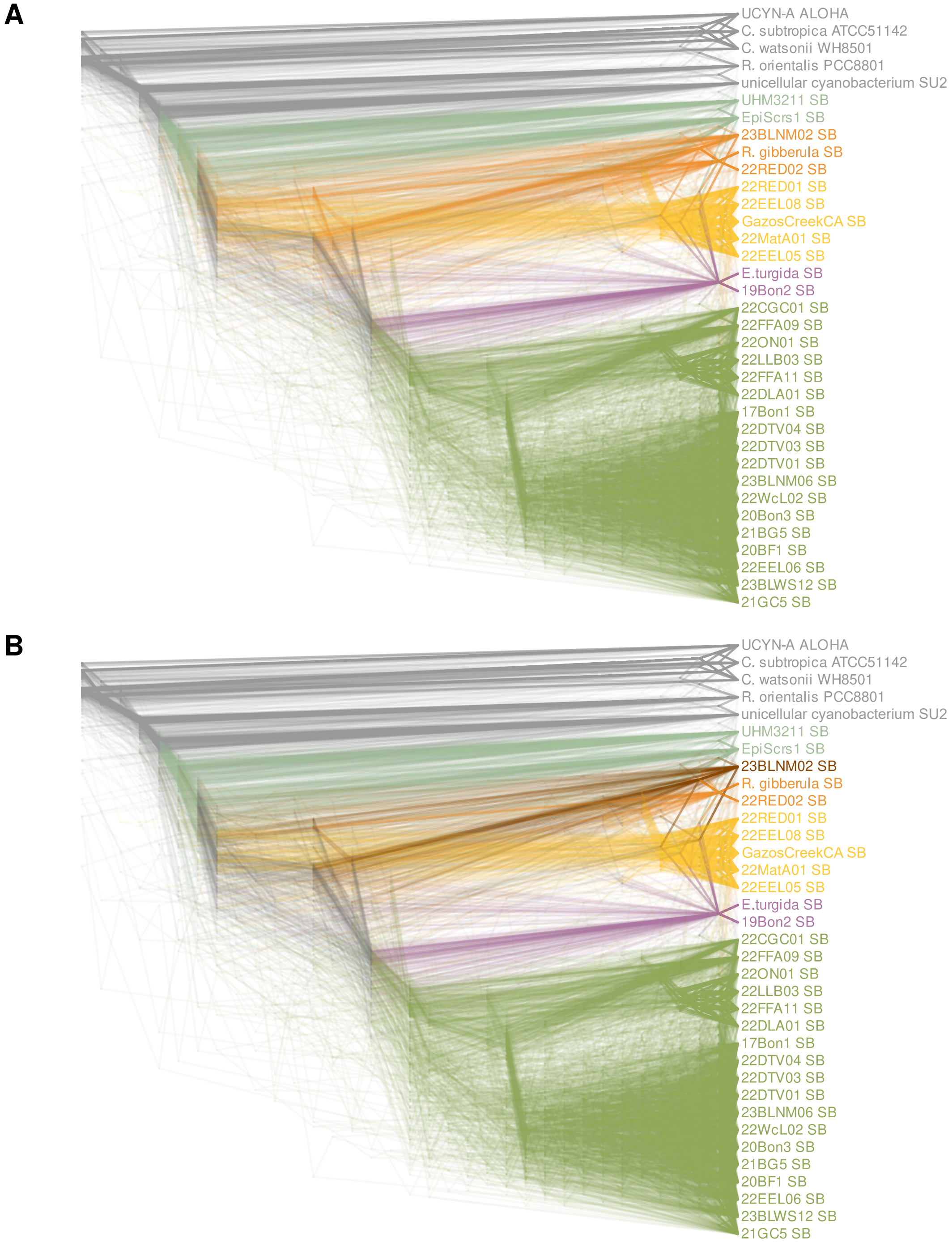


Figure S3: Cloudogram of 398 gene trees that were also used to generate the species tree presented in Figure 1 of the main text with branches colored according to their group. In A, 23BLNM02 SB is colored the same as the other two SBs from *R. gibberula*-like hosts. In B, 23BLNM02 SB is shown in a darker color to exemplify its varied placement in the gene trees. For visualization purposes in this figure, gene trees were mid-point rooted.

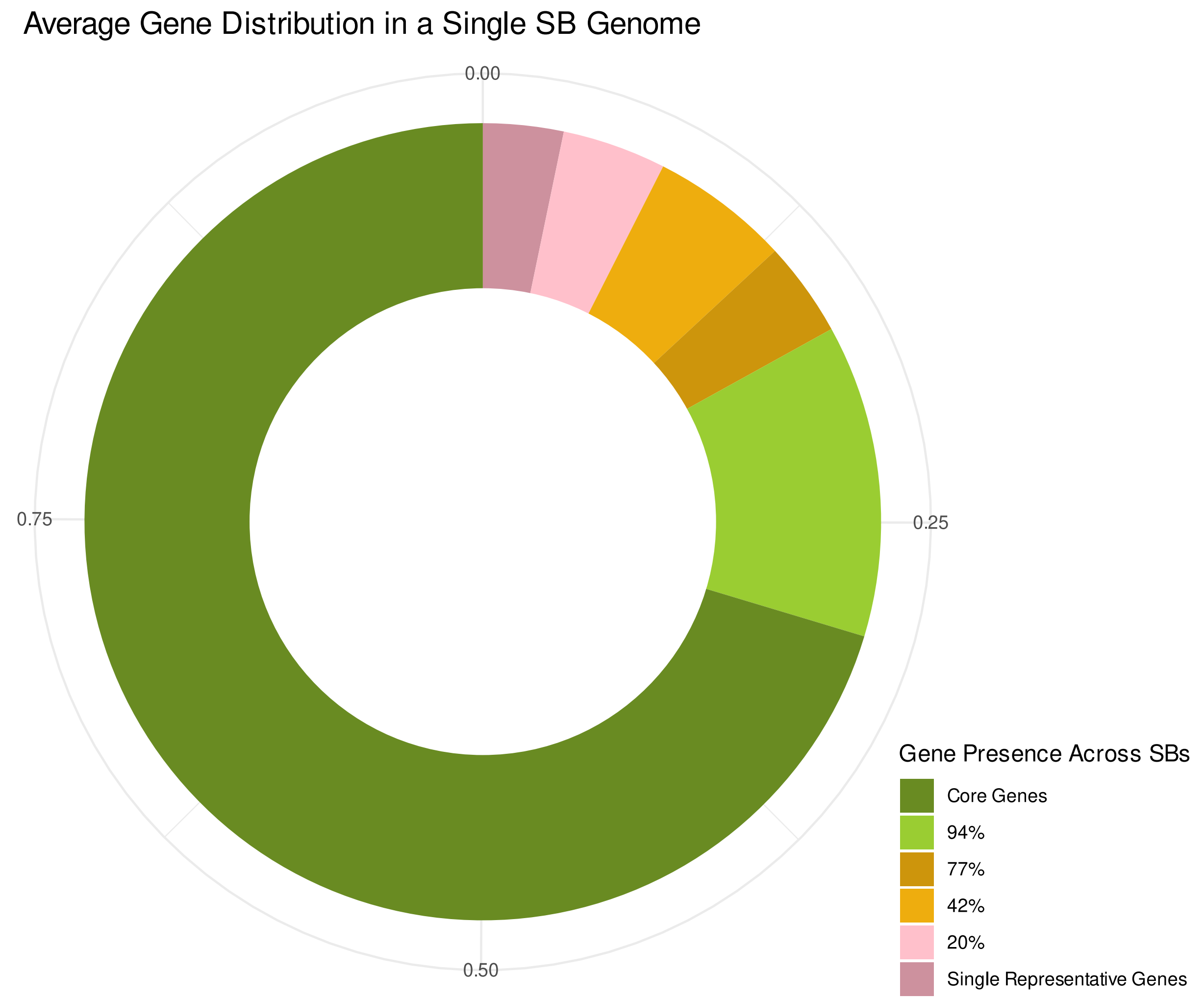


Figure S4: Gene content of the average SB genome, colored by how many SBs share the genes. Figure generated by the R package micropan.

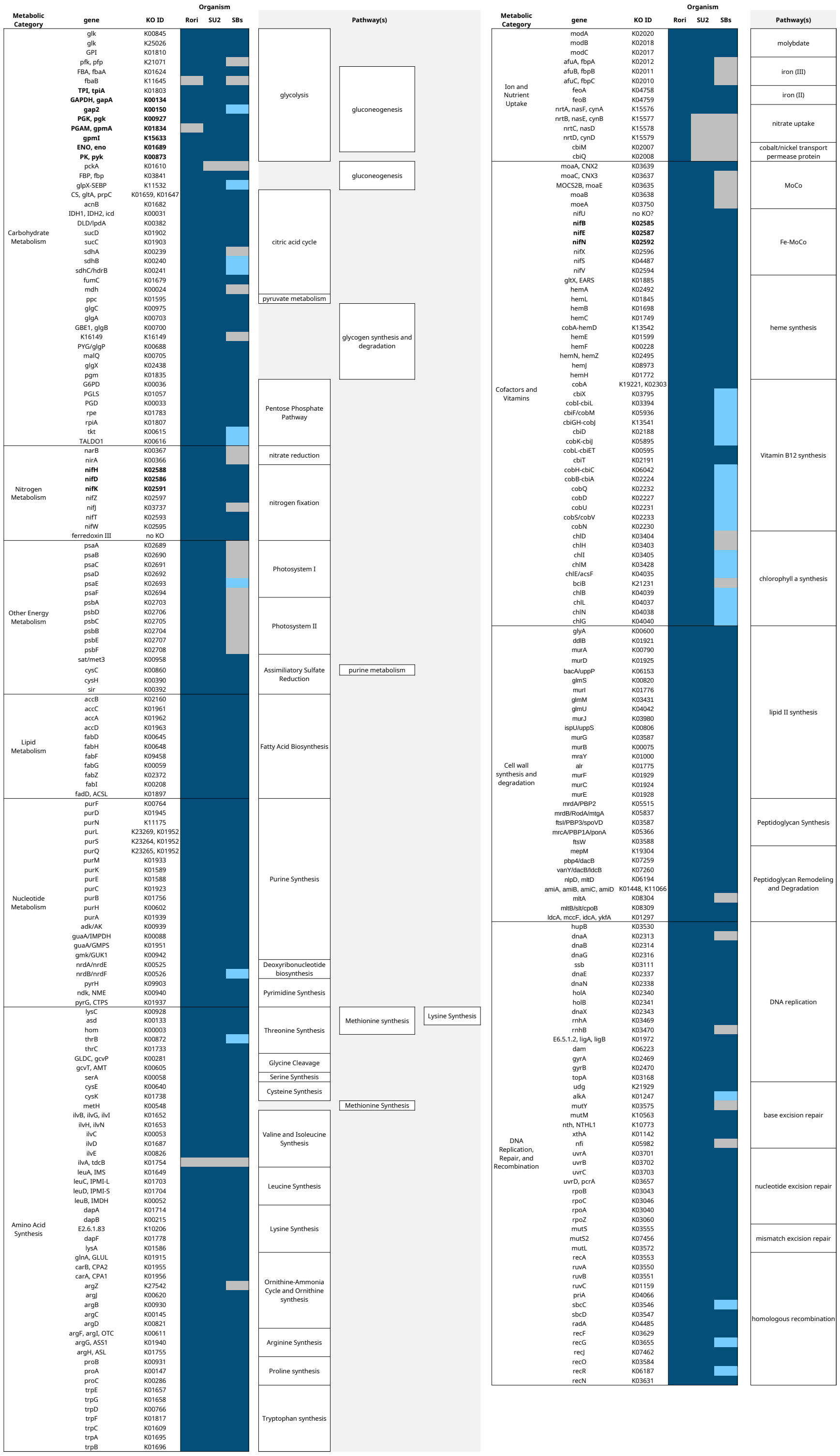

Figure S5: Presence and absence of genes for selected metabolic pathways in the SB core genome and two closely related free-living cyanobacteria. For all three categories, if the gene was not initially found by searching gene name or KO ID, blastp and/or tblastn searches were performed to confirm whether the gene was truly absent. Rori = *R. orientalis* PCC 8801; SU2 = unicellular cyanobacterium SU2; SBs = SB core genome; dark blue = present, present in all SBs; light blue = present in some SBs; grey = absent, absent in all SBs.

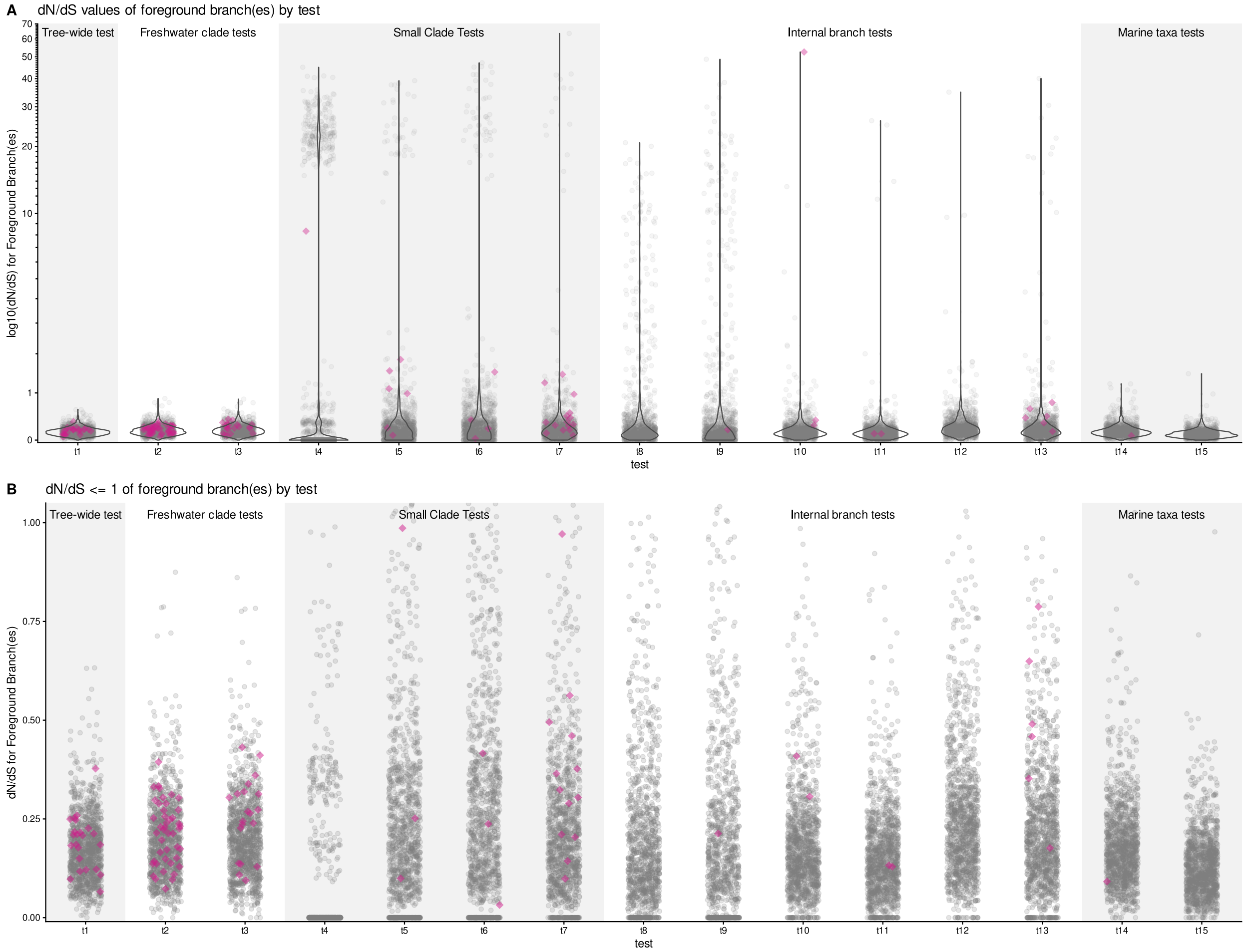


Figure S6: Distribution of dN/dS results for each test. A) all results and B) only dN/dS values less than 1. For both, significant LRTs for positive selection after correction are shown as pink diamonds. Note that A is shown on a log scale.
